# Paradise under threat: the successful invasion of the freshwater shrimp *Neocaridina davidi* and its parasites on La Réunion Island

**DOI:** 10.1101/2024.08.23.609356

**Authors:** Sebastian Prati, Laetitia Faivre, Valentin de Mazancourt, Bernd Sures, Pierre Valade

**Author notes:** Corresponding author: Sebastian Prati.

## Abstract

Invasive species are one of the major threats to global biodiversity, particularly in hotspots of endemic species, such as oceanic insular ecosystems. Tropical and subtropical islands are particularly vulnerable to biological invasions, with freshwater ecosystems expected to experience the most severe impacts. This is especially true if the insular freshwater fauna comprises a large set of near-threatened to critically endangered species.

This study assesses the establishment of the pet-traded shrimp *Neocaridina davidi* and its parasites on the island of la Réunion, a biodiversity hotspot within the Malagasy region. We confirm the establishment of feral *N. davidi* in two river catchments. Moreover, we confirm the presence of the microsporidian *Ecytonucleospora hepatopenaei*, an economically and ecologically relevant parasite that might impact a wide range of native species, including those living in brackish and marine environments. Three other microsporidians, two of which are unknown, have also been detected for the first time in *N. davidi*, raising concerns over possible spillover-spillback events. Therefore, we recommend timely actions to curb the spread of *N. davidi* and its parasites in an effort to preserve the already endangered native freshwater fauna.

## 1 Introduction

Due to their negative impact on recipient environments, invasive species are increasingly becoming one of the major threats to global biodiversity (Simberloff et al. 2013; Gallardo et al. 2016; Pyšek et al. 2020). This is especially true for hotspots of endemic species, such as oceanic insular ecosystems, where the impact of invasive species is disproportionally higher than that experienced on continents, mostly due to simplified food webs without top predators and small population sizes (Velmurugan 2008, Keppel et al. 2014, Russell et al. 2017, Lenzner et al. 2020). Tropical and subtropical islands are particularly vulnerable to biological invasions due to food-webs erosion bolstered by increased anthropogenic pressures and the introduction of non-native and eventually invasive species (Velmurugan 2008, Barlow et al. 2018, Moser et al. 2018). The largest impact of non-native species on oceanic islands is expected in terrestrial and freshwater ecosystems (Lenzner et al. 2020). Here, introduced non-native species often lead to strong predator-prey dynamics and competition, rapidly reducing the abundance of native species or leading to local extinctions (Russell et al. 2017). Freshwater ecosystems are generally more prone to experience biological invasion than their terrestrial counterpart due to habitat degradation and a soaring number of introduced non-native species linked to biological control, recreational angling, aquaculture, or pet trade (Cambray 2003, Savini et al. 2010, Patoka et al. 2018, Bernery et al. 2022). Even if introduced non-native species fail to establish, they might still negatively affect the recipient ecosystems by transmitting pathogens that can persist in the system even after the original host’s disappearance (Simberloff et al. 2013).

The oceanic island of La Réunion (French overseas territory), a biodiversity hotspot within the Malagasy region, is increasingly threatened by invasive species, including parasites introduced alongside their non-native hosts (Sasal et al. 2008, Soubeyran et al. 2015, Fenouillas et al. 2021). While the impact of invasive species on the island terrestrial ecosystem is relatively well-studied, information is lacking for the freshwater counterpart. Several of the 29 native fish and the vast majority of the 10 decapod species present on the island are considered locally near-threatened to critically endangered (Keith et al. 2006, Fricke et al. 2009, IUCN 2010, Froese and Pauly 2024). A situation that may worsen with the introduction of non-native species. Currently, successfully introduced non-native species on the island number eight fish and three decapods (Keith et al. 2006, Fricke et al. 2009, IUCN 2010, Leoville 2012, Froese and Pauly 2024). However, observations of additional species, mostly from the pet trade, suggest that the number of established non-native species is likely to increase. Several of the successfully introduced species on the island, e.g., *Amatitlania nigrofasciata*, *Cherax quadricarinatus*, *Poecilia reticuala*, and *Xiphophorus hellerii*, have also established feral populations worldwide mostly due to pet-trade releases (Magalhães and Jacobi 2017, Seebens et al. 2017, Baudry et al. 2020). Among pet-traded crustaceans not yet confirmed to be established on the island, and thus not included in the aforementioned list, is the Atyid shrimp, *Neocaridina davidi*. The shrimp was presumably observed on the island in 2017 by an employee of Hydrô Réunion, who signaled it to GEIR (Group Espèces Invasives de La Réunion). Volunteers of the latter, a group constituted by associations and professional and local institutions, promptly engaged in an eradication action as reported by a local newspaper (PM 2017). Since then, the trade/keeping of this shrimp has been banned on the island, but the feral population’s whereabouts remain unknown.

*Neocardina davidi*, a shrimp of Southeast Asian origins widely available in the global pet trade, displays high environmental plasticity and fecundity, which allow them to rapidly adapt to local conditions becoming invasive (Klotz et al. 2013, Prati et al. 2024). Established feral populations of *N. davidi* have been reported worldwide, including in Canada, Germany, Hungary, Israel, Japan, Poland, Slovakia, and the USA (Klotz et al. 2013, Jabłońska et al. 2018, deBruyn 2019, Levitt-Barmats et al. 2019, Weiperth et al. 2019, Prati et al. 2024). The ecological impacts of introduced *N. davidi* include the replacement of native shrimps with similar ecological niches (Onuki and Fuke 2022), alteration of meiofaunal assemblages (Weber and Traunspurger 2016), changes in leaf-litter breakdown in invaded areas (Schoolmann and Arndt 2017), and possible parasite exchange with native and invasive species (Prati et al. 2024).

*Neocaridina davidi* hosts a wide range of commensals and parasites (Ohtaka et al. 2012, Liao et al. 2018, Bauer et al. 2021, Maciaszek et al. 2023, Prati et al. 2024), some of which have been co-introduced outside their native range (Niwa and Ohtaka 2006, Maciaszek et al. 2021, Kakui and Komai 2022, Prati et al. 2024). Among them is also the ecologically and economically relevant parasite *Ecytonucleospora* (*=Enterocytozoon*) *hepatopenai* (EHP) (Wang et al. 2023), which has first been molecularly detected in a feral population of *N. davidi* inhabiting a German stream (Schneider et al. 2022). This parasite was later detected both molecularly and histologically in other European feral populations and animals sold in the pet trade (Prati et al. 2024). EHP is a host generalist parasite known to infect cultured Penaeid shrimps in coastal and marine environments (Chaijarasphong et al. 2021) but also a wide range of freshwater, brackish, and marine invertebrates, including the freshwater prawn *Macrobrachium rosenbergii* and insects such as dragonflies (Karthikeyan and Sudhakaran 2020, Krishnan et al. 2021, Jang et al. 2022, Munkongwongsiri et al. 2022, Dewangan et al. 2023, Wan Sajiri et al. 2023). However, EHP’s infectivity is reduced in low-salinity environments (Aranguren Caro et al. 2021).

Native freshwater decapods, most of which are considered vulnerable, comprise several amphidromous species belonging to the genera *Atyoida*, *Caridina*, *Macrobrachium*, and *Palemon* (Keith et al. 2006). These might be impacted by introduced *N. davidi* and their parasites. If present, infected *N. davidi* individuals might spread pathogens to native species. In the case of EHP, the parasite might also indirectly impact other macroinvertebrates, including those living in brackish and marine environments. Therefore, this study aims to confirm the presence of feral *N. davidi* on the island and conduct a parasitological screening.

## 2 Materials & methods

### 2.1 Sampling sites

Shrimps were sampled on the 15th of April, 2024, in three sites along two river catchments with perennial water on the island’s eastern side: the Mat (n=2) and the Saint-Jean (n=1). The Mat, with 145 km², is the largest catchment on the island. Its water sources, fed by regular rainfall, originate from the volcanic cirque of Salazie and flow eastwards toward the Indian Ocean. In the lower part of the river, a seemingly large population of shrimps, presumably *N. davidi*, was observed for the first time in 2017 by a group of citizen conservationists (PM 2017).

Successively, riverine ecological monitoring campaigns conducted by OCEA consult revealed a widespread presence of shrimps in the Mat catchment. These monitoring campaigns also included other areas, revealing the shrimp’s presence in the lower part of the nearby Saint-Jean catchment (Figure 1). We, therefore, collected shrimps in the upper part of the Mat River from a right-side creek outflowing from the Mare à Poule d’Eau, a pond located in Salazie, from the Bras Citronnier, a left-side tributary of the Mat River located further downstream, and from the Ravine Sèche, a right-side tributary of the Saint-Jean River (Figure 1 and 2, Table 1). At the Ravine Sèche site, presumed *N. davidi* co-occurred with the native *C. typus*.

**Figure 1.**
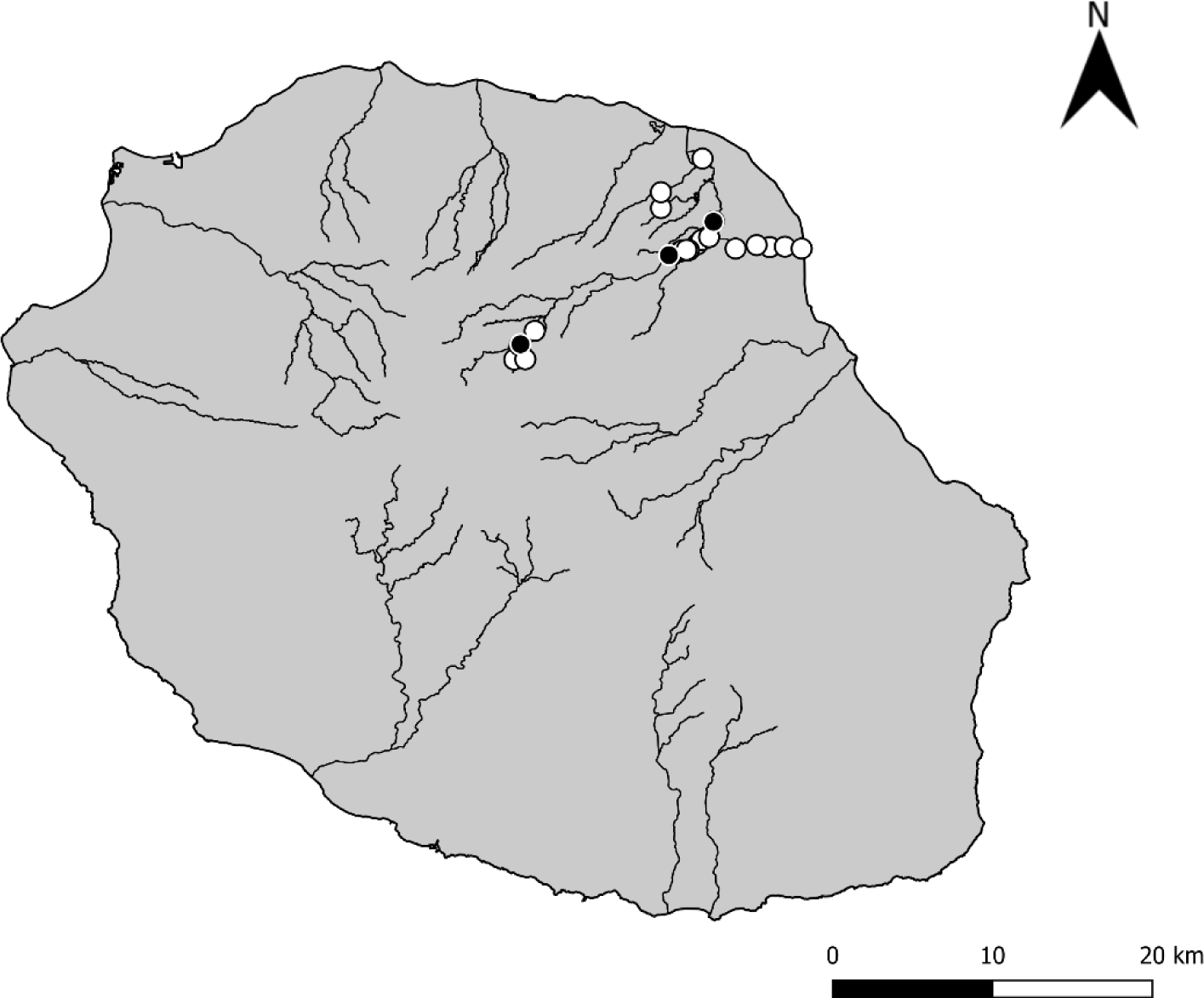
Map of the sampling areas where *Neocaridina davidi* individuals were collected. Black dots indicate sites where shrimp were collected in the present study, and white dots indicate additional sites where *N. davidi* was observed during monitoring campaigns conducted by OCEA consult. The map was generated using QGIS.

**Figure 2.**
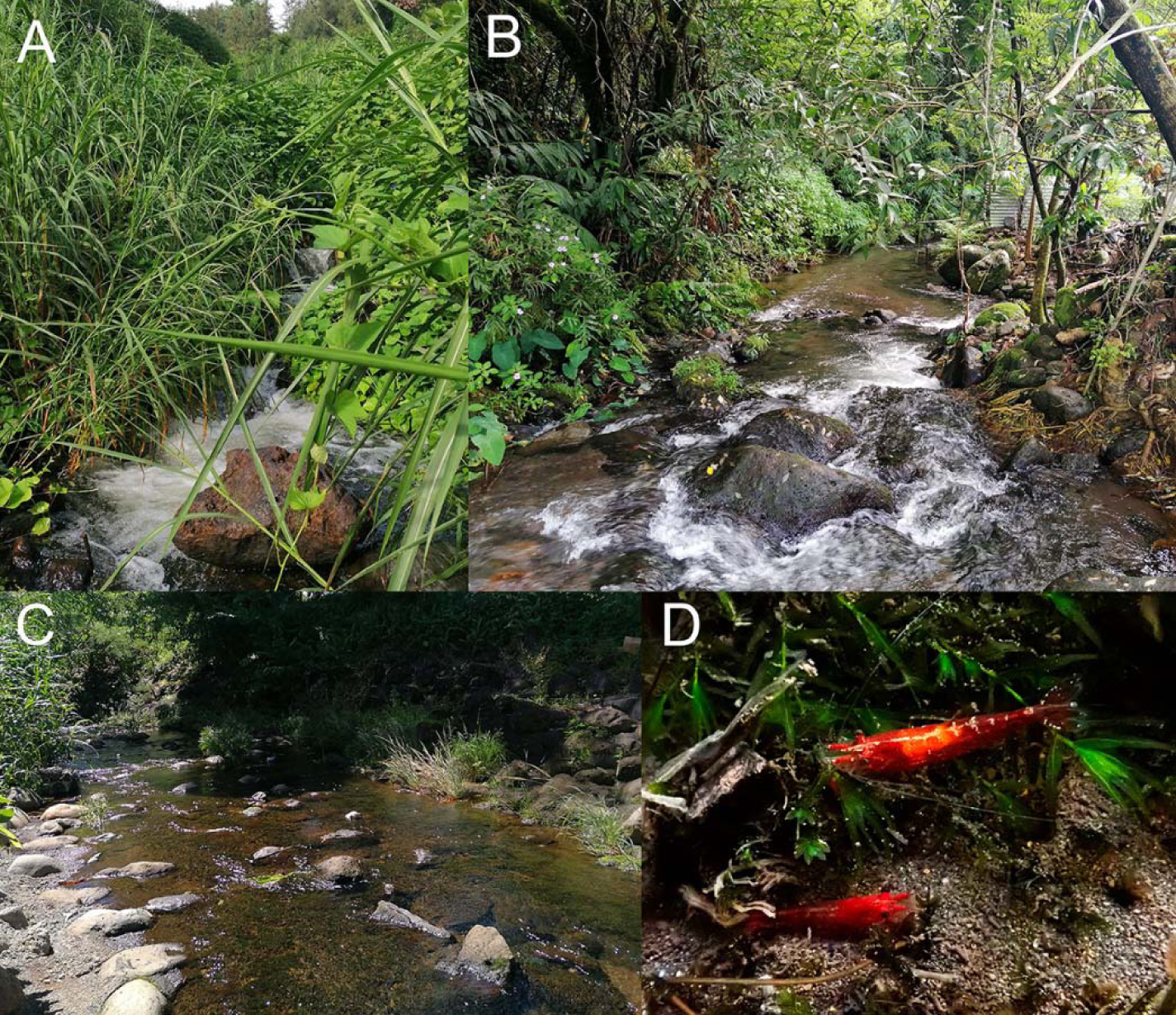
Sites in which *Neocaridina davidi* were collected: **A** Creek near Salazie, **B** Bras Citronnier, and **C** Ravine Sèche, and **D** feral *Neocaridina davidi* photographed underwater.

**Table 1.** Sampling sites in which *N. davidi* were collected, including elevation, temperature, and conductivity measured at the collection date (15/04/2024).

| Catchment | Sampling sites | Coordinates | Elevation m.s.l. | Temperature [°C] | Conductivity [µs/cm] |
| --- | --- | --- | --- | --- | --- |
| Mat | Creek near Salazie | -21.0418, 55.5295 | 526 | 24.5 | 268 |
|  | Bras Citronnier | -20.9881, 55.6191 | 171 | 24.5 | 132 |
| Saint-Jean | Ravine Sèche | -20.9680, 55.6460 | 120 | 26.2 | 76 |

### 2.2 Sampling collection and processing

Shrimps were collected using hand nets and immediately preserved in 96% ethanol. All individuals were morphologically and molecularly identified, measured, dissected in the laboratory, and visually screened for internal parasites. The morphological identification of presumed *N. davidi*, followed (Englund and Cai 1999, Klotz et al. 2013). However, its taxonomy remains somewhat unclear, and the names *N. davidi* (syn. *N. denticulata sinensis* and *N. heteropoda*) and *N. denticulata* are often used interchangeably in literature (e.g., Levitt-Barmats et al. 2019; Onuki and Fuke 2022). Therefore, the distinction between *N. davidi* and *N. denticulata* relied on the sexual dimorphism in the third pereiopods and the distinctly shorter appendix interna of the male second pleopods, as suggested by Shih et al. (2024). A small portion of shrimp, including muscles and hepatopancreatic tissue, was used for molecular analyses of hosts and microsporidian parasites. Epibionts were not analyzed as their preservation in ethanol is generally poor.

### 2.3 Molecular analyses

DNA was isolated from hosts using a modified salt precipitation protocol described by Grabner et al. (2015). Shrimps were identified using the universal eukaryotic primers LCO1490 (5′-GGTCAACAAATCATAAAGATATTGG-3′) and HCO2198 (5′ TAAACTTCAGGGTGACCAAAAAATCA-3′) (Folmer et al. 1994) targeting the COI region and microsporidians with the universal microsporidian primer V1F (5′-CACCAGGTTGATTCTGCCTGAC-3′) (Zhu et al. 1993) and 1342R (5′-ACGGGCGGTGTGTACAAAGAACAG-3′) (McClymont et al. 2005) targeting the small subunit ribosomal RNA gene (SSU rRNA). PCR reaction for shrimps consisted of 20 μL assay with 10 μL of Dream-TaqTM Hot Start Green PCR Master Mix (Thermo Fisher Scientific, Waltham, MA, USA), 1.6 μL (5 uM) of each primer, 4.8 μL of nuclease-free water and 2 uL of DNA template per reaction. Those for microsporidans consisted of 20 μL composed of 10 μL of 2x AccuStart II PCR ToughMix (Quantabio), 1 μL of each primer (0.5 μM), 0.35 μL of 50x GelTrack Loading Dye (Quantabio), 6.65 μL MilliQ water and 1 μL of DNA template. The PCR setting used for the primer pair LCO1490-HCO2198 followed those employed by Prati et al. 2024), while those used for the primer pair V1F-1342R were the following: initial denaturation for 5 min at 94 °C, followed by 35 cycles of denaturation for 40 s at 94 °C, 40 s of annealing at 60 °C and 55 s of elongation at 68 °C, and a final elongation step for 5 min at 68 °C. PCR products were sent unpurified to Microsynth Seqlab (Germany) for Sanger sequencing.

Raw sequences were quality-checked and edited using Geneious Prime v2024.0.5 (Biomatters, Ltd., New Zealand) and then compared against GenBank records using the blastn algorithm (https://blast.ncbi.nlm.nih.gov). Shrimps and parasite sequences were separately aligned using the MAFFT v7.490 algorithm with standard settings (Katoh et al. 2019) and Maximum likelihood phylogenetic trees (1000 replicate bootstrap values) were produced in IQ-Tree 2.2.2.6 (Minh et al. 2020). The substitution models chosen based on the Bayesian Information Criterion were TPM2u+F+I for shrimps and TIM3+F+G4 for microsporidians, and the outgroups *Neocaridina palmata* (NCBI GenBank accession number MN701612) and *Triwangia caridinae* (JQ268567). Sequences generated in this study were submitted to NCBI GenBank (accession number PQ069719 for N. davidi and PQ096043-47 for microsporidians).

### 2.4 Statistical analyses

Differences in the carapax length among *N. davidi* collected at different sampling sites were investigated with linear regression, accounting for size differences related to sex. The model included carapax length as the dependent variable and sampling sites and sex (females and males) as interacting independent variables. The results were summarized using the Anova type 3 function from the r package car (Fox and Weisberg 2018). The prevalence of microsporidian parasites was too low for direct comparisons between shrimp populations or shrimp sex. However, we used the exact binomial probability estimation with a 95% confidence interval to estimate the infection rate of background populations at each site using the r package exactici (Fay 2023). Statistical and descriptive analyses were performed with the open-source software R (version 4.4.1, R Core Team 2024) via the RStudio GUI (version 2024.04.2, RStudio Inc.).

## 3 Results

A total of 120 presumed *N. davidi* shrimp were collected. The sexual dimorphism in the third pereiopods and appendix interna of the male second pleopods was consistent with that of *N. davidi*. Moreover, the shrimps showed a bright red coloration typically displayed in the pet-traded *N. davidi* sold under the commercial names red cherry and red fire (Figure 2 D). Morphological identification was corroborated by molecular identification. Irrespective of sampling location, a single *N. davidi* haplotype based on COI sequences was detected. This showed 100% similarity and 97.04-100% coverage with the haplotype Ndh2 (see Prati et al. 2024) of feral and pet-traded individuals previously found in Brazil (PP718677), Canada (MG319788), Germany, Hungary, Poland, and Slovakia (MG816764-65 and OR610863), and wild individuals from Taiwan (MG734262) (Figure 3).

**Figure 3.**
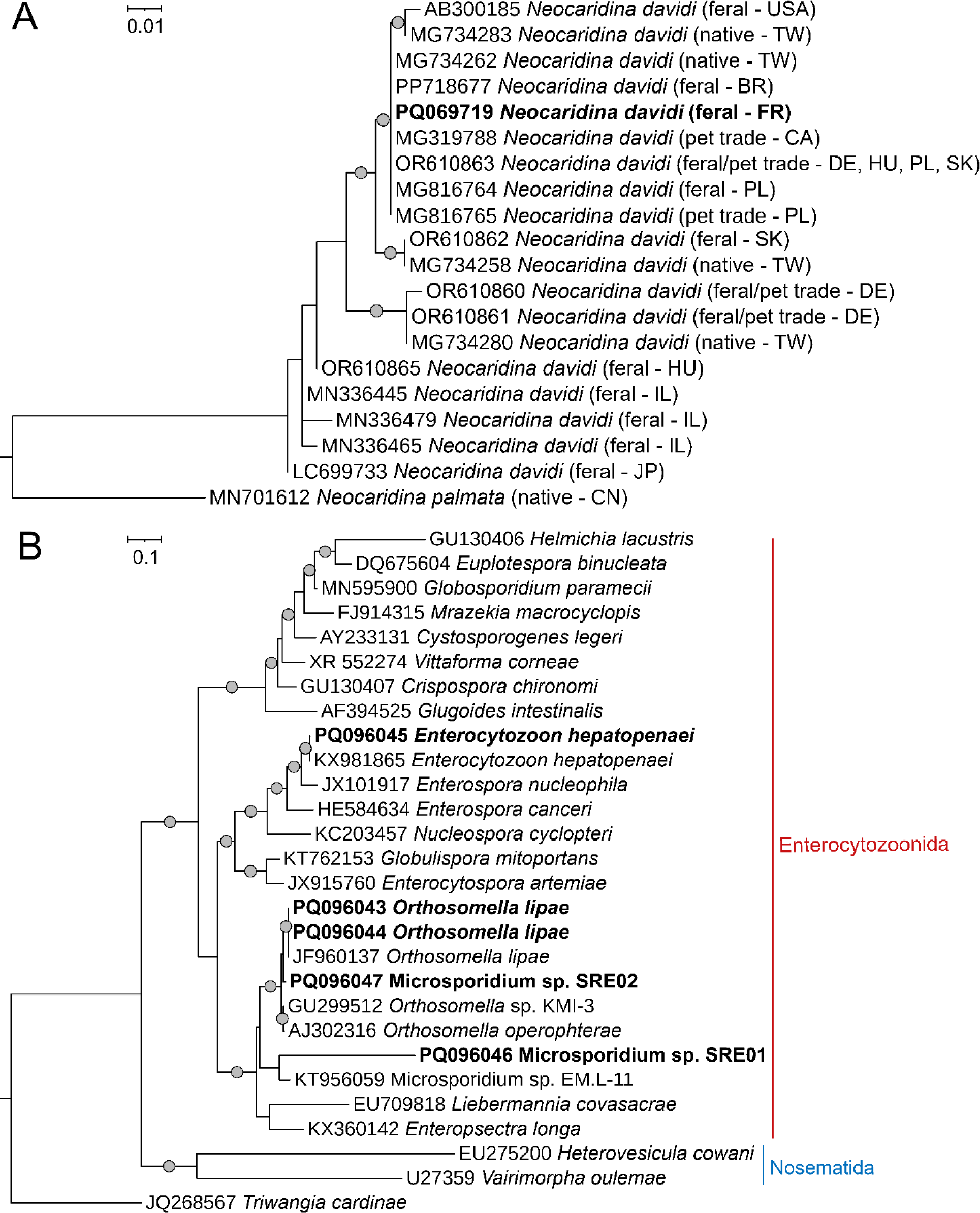
**A** Maximum likelihood phylogenetic tree of feral *Neocaridina davidi* and **B** microsporidians identified in this study. Grey dots represent bootstrap support values above 90%. Sequences obtained in this study are indicated in bold. The substitution models used are TPM2u+F+I for shrimps and TIM3+F+G4 for microsporidians, and the outgroups are represented by *Neocaridina palmata* and *Triwangia caridinae*. The names and circumscriptions of microsporidian clades follow Bojko et al. (2022).

Among the collected *N. davidi*, 24 were females (four of whom were ovigerous), 56 males, and 40 immatures. Overall, the size of sexed *N. davidi* varied greatly between genders (*F* (1, 74) = 5.1, *p* = 0.026), with females having a larger carapax length than males and between sampling sites (*F* (2, 74) = 16.14, *p* <0.001) mostly due to the large size of individuals of both sexes collected at Ravine Sèche. However, such differences were also influenced by differing sex ratios across sampling sites (*F* (2, 74) = 4.31, *p* = 0.017). Namely, the highest proportion of females was found in the Bras Citronnier (36.36%) and the lowest at the ravine Ravine Sèche site (26.47%). The overall prevalence of microsporidians in the shrimp investigated was 4.16% and between 1.36 and 9.46% for background populations. Four isolates were detected, three of which were found in the Bras Citronnier population (Table 2). Immature *N. davidi* individuals had the highest diversity of microsporidians, albeit with low prevalences. No females but a single male was infected (Table 2).

**Table 2.** Prevalence of microsporidians in *Neocaridina davidi* for every sampling site (All = all, F = females, M = males, and I = immatures individuals), with a 95% confidence interval for background population prevalence based on binomial probability estimation. Note that the 95% confidence interval is calculated for the overall prevalence and that of each microsporidium based on all individuals (n=40) collected at each sampling site.

| Sampling sites | N | Sex | Size | Prevalence in % | 95% CI in % | Microsporidians |
| --- | --- | --- | --- | --- | --- | --- |
| Creek near Salazie | 40 | All | 3.19 ± 0.91 | 5 (2/40) | 0.61-16.92 | All |
|  | 7 | F | 4.23 ± 0.61 | 0 (0/7) | - | - |
|  | 17 | M | 3.66 ± 0.26 | 0 (0/17) | - | - |
|  | 16 | I | 2.23 ± 0.44 | 12.5 (2/14) | 0.61-16.92 | *Orthosomella lipae* |
| Bras Citronnier | 40 | All | 3.07 ± 0.88 | 7.5% (3/40) | 1.57-20.39 | All |
|  | 8 | F | 4.05 ± 0.48 | 0 (0/8) | - | - |
|  | 14 | M | 3.57 ± 0.29 | 7.14 (1/14) | 0.06-13.16 | Microsporidium sp. SRE01 |
|  | 18 | I | 2.24 ± 0.46 | 5.56 (1/18) | 0.06-13.16 | *Ecytonucleospora hepatopenaei* |
|  |  |  |  | 5.56 (1/18) | 0.06-13.16 | Microsporidium sp. SRE02 |
| Ravine Sèche | 40 | All |  | 0 (0/40) | 0.00-8.81 | All |
|  | 9 | F | 5.27 ± 0.77 | 0 (0/9) | - | - |
|  | 25 | M | 4.04 ± 0.50 | 0 (0/25) | - | - |
|  | 6 | I | 2.62 ± 0.36 | 0 (0/6) | - | - |

*Neocaridina davidi* were infected with EHP isolated from the prawn *Penaeus vannamei* (99.9% similarity and 100% coverage to KX981865), *Orthosomella lipae* isolated from the coleopteran *Liophloeus lentus* (100% similarity and 99.5% coverage to JF960137), and two undescribed microsporidian isolates here named Microsporidum sp. SRE01 and Microsporium sp. SRE02. The most similar isolate to Microsporidum sp. SRE01 was Microsporidium sp. EM.L-11 isolated from the amphipod *Eulimnogammarus verrucosus* (87% similarity and 84.2% coverage to KT956059), and the closest described species was *Enteropsectra longa* isolated from the nematode *Oscheius* sp. (85.3% similarity and 84.5% coverage, to KX36014). The most similar isolate to Microsporidum sp. SRE02 was *Orthosomella* sp. KMI-3 isolated from the moth *Conistra vaccinii* (98.2% similarity and 100% coverage to GU299512), and the closest described species was *Orthosomella operophterae* isolated from the moth *Operophtera brumata* (97.9% similarity and 100% coverage to AJ302316). All detected microsporidians belonged to the *Enterocytozoonida* clade sensu Bojko et al. (2022).

## 4 Discussion

Insular freshwater ecosystems are increasingly threatened by invasive species, including those of pet-traded origin (Velmurugan 2008, Moser et al. 2018, Baudry et al. 2020, Lenzner et al. 2020). The establishment of *N. davidi* and associated parasites in the Mat and the Saint-Jean catchments on the island of Réunion is one such example. Commercially successful pets like *N. davidi* are inexpensive, widely available, prolific, and adaptable to a wide range of environmental variables. Features that favor their establishment in novel locations (Lipták and Vitázková 2015, Bláha et al. 2022, Prati et al. 2024). Their commercial success depends on mass production, which creates an ideal environment for the proliferation of pathogens, including parasites (Maciaszek et al. 2023, Prati et al. 2024). Due to a lack of stringent biosecurity measures along the supply chain, these might end up in private aquaria (Patoka et al. 2016, 2018, Maciaszek et al. 2021, Prati et al. 2024). Consequently, releasing ornamental pets and associated parasites into novel environments might pose severe risks to native biota (Sures 2011, Svoboda et al. 2017, Patoka et al. 2018). Therefore, deepening the knowledge of invading hosts and parasites is paramount, especially in insular systems.

Feral *N. davidi* populations have been present in Europe since 2003 when a population established itself in Poland (Jabłońska et al. 2018). Afterward, more established populations were discovered in Germany, Hungary, and Slovakia (Klotz et al. 2013; Weiperth et al. 2019; Prati et al. 2024). The present study expands this list to two river catchments in the French overseas territory of la Réunion, increasing the number of confirmed established non-native decapods on the island to three. The presence of *N. davidi* haplotype Ndh2 on the island is unsurprising, as this is the most common haplotype found in feral and pet-traded individuals (Prati et al. 2024). The occurrence of a single haplotype suggests that the collected individuals stem from a single source population, which might have been translocated across river catchments. This lack of genetic diversity compared to other feral populations likely reflects the remoteness of the island and the difficult access to a larger variety of pet-traded *N. davidi*.

Surprisingly, all shrimps displayed a bright red coloration, similar to that displayed by a feral population in Canada (deBruyn 2019). This is rather uncommon, as feral *N. davidi*, similarly to other crustaceans (Duarte et al. 2017), rely on camouflage to avoid predation, thus reverting to nearly translucent to brown coloration typically found in wild populations (Englund and Cai 1999, Levitt-Barmats et al. 2019). Accordingly, flashy colorations, which are more accentuated in females, likely result in increased predation pressures by visual feeders such as fish (Prati et al. 2024). Indeed, intense predation by both native and non-native species upon feral *N. davidi* has been recorded in a German stream (Schoolmann and Arndt 2018). This might not be the case on the island’s freshwater ecosystems, where top predators are lacking. On the island of la Réunion, the only top predator in freshwater ecosystems is the non-native *Oncorhynchus mykiss,* which is found in the colder water of rivers located at higher elevations (Keith 2002, Keith et al. 2006).

Native fish species, e.g., *Anguilla* spp., *Awaous commersoni*, *Cotylopus acutipinnins*, *Eleotris* sp., and *Kulia rupestris* might potentially feed on *N. davidi* and are present in the lower and middle stretches of the two investigated catchments (Biotope 2021). However, their abundance is dwindling, with all of the abovementioned species being classified as near-threatened, vulnerable, or critically endangered (IUCN 2010). Moreover, the presence of impoundments along the rivers might further limit their occurrence and movement. Thus, the predation pressure asserted by native species on *N. davidi* is nearly nonexistent. Predation from non-native species seems also negligible. Non-native fish species such as *P. reticulata* are commonly found co-occurring with *N. davidi*. However, their small mouth allows only for a short window in which the predation of, i.e., newborn shrimp, can occur.

This lack of predation is also evidenced by the sex ratio of collected *N. davidi* individuals. In *N. davidi*, the sex ratio is temperature dependent, with females accounting for about 20% of the population at 26°C and just over 50% at 23°C under laboratory conditions (Serezli et al. 2017). Females, which are larger and more colorful than males, are more likely to be targeted by visual predators like fish; therefore, in their presence, a lower proportion of females than what can be predicted at a certain water temperature is expected. Based on the water temperature at the sampling sites, the sex ratio among the collected individuals reflected what was expected in the absence of predators. Therefore, without effective top-down control, *N. davidi* will continue to proliferate undisturbed on the island rivers.

The further proliferation of *N. davidi* on the island’s rivers could potentially pose a severe threat to native decapods, particularly amphidromous shrimp with similar niches such as *Atyoida serrata*, *Caridina natalensis*, and *Caridina typus*. All of these species are classified as either near threatened or vulnerable (IUCN 2010). In the present study, only a few *C. typus* individuals were observed co-occurring with *N. davidi* at the Ravine Sèche site. This situation underscores the potential impact of *N. davidi* on native decapods, emphasizing the severity of the issue. Due to its high feeding rate, which is estimated to be over half of its body weight per day, *N. davidi* might outcompete and replace less efficient native crustaceans who rely on the same resources (Schoolmann & Arndt 2017; Onuki and Fuke 2022).

Trophic niche aside, *N. davidi* possesses a major competitive advantage over the native shrimps due to high reproduction rates and a direct development unfolding entirely in freshwater (Tropea and Greco 2015). Females of *N. davidi* produce up to 60 eggs that hatch within 16-19 days at 25°C and may develop new ones right after the previous batch hatched (Pantaleão et al. 2017, Schoolmann and Arndt 2018). Newborns can reach sexual maturity within 3 months (Pantaleão et al. 2017). On the other hand, all native species are amphidromous, meaning their freshly hatched larvae must reach marine environments to complete their development. However, anthropogenic interference on the island, such as river discharge alteration and occasional dewatering of river mouths, can considerably impact larvae drift toward the ocean (Hoarau et al. 2019). Moreover, the adult stages of native species are considered a local delicacy and subject to overfishing (Hoarau et al. 2019). Another aspect that may impact native species and should be considered is the possible transmission introduction of parasites alongside *N. davidi*.

The investigated *N. davidi* were infected with four different microsporidians, three of which were recorded for the first time in this host, increasing the number of its endoparasites to seven. Among them is the microsporidian parasite EHP, which was previously reported from pet traded and feral populations (Schneider et al. 2022, Prati et al. 2024). This parasite, which is tolerant to a wider range of salinity (up to 40 ppt) has been reported from a variety of invertebrate hosts (Karthikeyan and Sudhakaran 2020, Aranguren Caro et al. 2021, Krishnan et al. 2021, Munkongwongsiri et al. 2022, Wan Sajiri et al. 2023). Its presence on the island is concerning, as the parasite might spread to native biota via predation, e.g., in the case of dragonflies, or via spore ingestions in shrimps or other macroinvertebrates feeding on contaminated organic matter (Dewangan et al. 2023). Moreover, spores might be transported by water flow and end up in brackish and marine environments, where they are more infective (Aranguren Caro et al. 2021). *Neocaridina davidi*’s tolerance to salinity is unknown, but the recent finding in Brazil of a single individual in seawater (Bochini et al. 2024) opens novel possibilities for the spread of this invasive species and its parasites. Fortunately, the observed prevalence of EHP remains low, likely due to the lotic nature of the environments inhabited by *N. davidi,* which may contribute to a dilution of infective spores. Nevertheless, the current background population prevalence might be as high as 13.6%. Thus, the situation should be closely monitored, as the consequences of a possible outbreak among native species are unpredictable.

Parasites that are transmitted from introduced to native host species may have greater pathogenic effects in native hosts as they lack a coevolutionary history (Mastitsky et al. 2010). However, non-native species might also acquire local parasites, amplifying or diluting their prevalence in native species (Strauss et al. 2012, Telfer and Bown 2012). This might be the case for the other three microsporidians detected in the present study. It is not known if these microsporidians are present in native species, but until now, they have never been detected in feral or pet-traded *N. davidi*, which suggests local origin. Members of the genus *Orthosomella*, to which two of our sequences belong, are parasites of insects, but undescribed isolates have been also detected in gastropods suggesting possible switching between hosts that share similar habitats (Ovcharenko et al. 2013). Unfortunately, the study of parasites co-introduced and acquired by introduced species and the consequences for local environments are often neglected in biological invasion studies. Therefore, we recommend timely actions to curb the spread of *N. davidi* and its parasites on the island in an effort to preserve the already endangered native freshwater fauna.

## Statements and Declarations

### Funding

This study was performed within the Collaborative Research Center (CRC) RESIST (A09) and funded by the Deutsche Forschungsgemeinschaft (DFG, German Research Foundation) – CRC 1439/1 – project number: 426547801. We acknowledge support from the Open Access Publication Fund of the University of Duisburg-Essen.

### Ethics and Permits

French authority waived ethical approval for this study under order No. 23-05/DEAL/SEB/UPEMA, authorizing Pierre Valade to capture and transport fish and macrocrustaceans for ecological purposes as part of his mission as head of the GEIR network.

### Author contributions

Sebastian Prati, Laetitia Faivre, Valentin de Mazancourt, Bernd Sures, and Pierre Valade conceived the study. Pierre Valade supervised the project. Sebastian Prati and Pierre Valade carried out the sampling. Sebastian Prati carried out laboratory analyses and data analyses. Sebastian Prati wrote the first draft of the manuscript, and all authors contributed critically to the final draft. All authors read and approved the final manuscript.

### Competing interests

The authors have no relevant financial or non-financial interests to disclose.

## Acknowledgments

The authors have no non-financial support to report.

## Data availability

The reported nucleotide sequence data are available in the GenBank database under the accession numbers 9719 for *N. davidi* and 6043-47 for microsporidians.

## Notes

### Competing Interest Statement

The authors have declared no competing interest.

